# Aligned and Conductive 3D Collagen Scaffolds for Skeletal Muscle Tissue Engineering

**DOI:** 10.1101/2020.04.18.048017

**Authors:** Ivan M. Basurto, Mark T. Mora, Gregg M. Gardner, George J. Christ, Steven R. Caliari

## Abstract

Skeletal muscle is characterized by its three-dimensional (3D) anisotropic architecture composed of highly aligned, organized, and electrically excitable muscle fibers that enable normal locomotion. Biomaterial-based tissue engineering approaches to repair skeletal muscle injuries are limited due to difficulties in combining 3D structural alignment (to guide cell/matrix organization) and electrical conductivity (to enable electrically excitable myotube assembly and maturation). In this work we successfully produced aligned and electrically conductive 3D collagen scaffolds using a freeze-drying approach. Conductive polypyrrole (PPy) microparticles were synthesized and directly mixed into a suspension of type I collagen and chondroitin sulfate followed by directional lyophilization. Scanning electron microscopy (SEM), energy-dispersive spectroscopy (EDS), and confocal microscopy analyses showed that directional solidification resulted in scaffolds with longitudinally aligned macropores (transverse plane: 155 ± 27 µm, longitudinal plane: 218 ± 49 µm) with homogeneously-distributed PPy content. Chronopotentiometry verified that PPy incorporation resulted in a five-fold increase in conductivity when compared to non-PPy containing collagen scaffolds without detrimentally affecting C2C12 mouse myoblast metabolic activity. Furthermore, the aligned scaffold microstructure provided contact guidance cues that directed myoblast growth and organization. Incorporation of PPy also promoted enhanced myotube formation and maturation as measured by myosin heavy chain (MHC) expression and number of nuclei per myotube. Together these data suggest that aligned and conductive 3D collagen scaffolds could be useful for skeletal muscle tissue engineering.

## 1. Introduction

Skeletal muscle tissue possesses an innate ability to heal from most injuries by employing an inflammatory response that results in a cycle of degeneration, remodeling, and repair [1,2]. However, the magnitude and multi-tissue nature (e.g., muscle, nerve, vasculature) of volumetric muscle loss (VML) injuries compromises the wound healing response and leads to a permanent loss of muscle mass and function, impacting both civilian and military populations [3–5]. In fact, musculoskeletal trauma constitutes 65% of injuries sustained in recent military conflicts and 92% of soldiers with muscle deformities are classified as VML patients [4,6]. VML-type injuries also account for a significant portion of civilian injuries including myopathies, surgical loss, and traumatic injuries such as car accidents, gunshot wounds, and compound fractures [7].

The inability to more effectively treat VML injuries in both military and civilian populations motivates the development of therapies that can more fully restore muscle form and function. Current approaches include the autologous transfer of healthy skeletal muscle from a donor site to the area of injury followed by extensive physical therapy [8]. However, this treatment method has proven both expensive and time-intensive, resulting in generally poor levels of functional improvement as well as associated donor site morbidity that limits the size of the injury that can be treated [9]. Further limitations to effective treatments include the considerable variability that exists between injuries including size, depth, and location as well as the heterogeneity of affected tissue types [10]. As a result, there is currently no effective treatment available to regenerate large areas of musculoskeletal tissue. In an effort to address this important unmet medical need research has focused on tissue engineering/regenerative medicine approaches, specifically the use of acellular extracellular matrix (ECM) scaffolds, seeded with myogenic cell sources for subsequent implantation [11–13]. Other approaches have employed rationally engineered biomaterial platforms that provide chemical and physical cues to direct cell survival and more accurately mimic healthy ECM microenvironments [14–17]. Numerous studies also combine biomaterials with isolated muscle cells as well as supporting cells including neurons, endothelial cells, and fibroblasts to augment the repair process [11,18–22]. While many of these strategies hold promise, they are still limited in their ability to drive functional, innervated, and vascularized muscle repair, based in part on their inability to mimic both the structural organization and the electrochemical excitability of native muscle.

Skeletal muscle development proceeds in a tightly regulated manner as myocytes fuse to form myotubes [23] and progressively organize into aligned myofibers, fascicles, and larger muscle fiber bundles [23,24]. During this process, a complex ECM is deposited surrounding muscle fibers (endomysium), muscle fascicles (perimysium), and the muscle body (epimysium) composed primarily of type I and III collagen [25]. The ECM also includes type IV and V collagen as well as proteoglycans such as decorin, biglycan, and laminin [25]. It is this anisotropic organization from the molecular to tissue scale that ultimately allows for effective force generation during muscle contraction [25]. However, in the case of myopathies and severe trauma like VML the ECM becomes disorganized and results in a permanent loss of muscle function [10], thus motivating the use of regenerative templates that direct anisotropic cell and ECM organization following injury. In addition to the highly organized structure of skeletal muscle ECM, bioelectrical stimuli in the cellular microenvironment can regulate cell fate decisions involved in embryonic development, myoblast differentiation, and tissue regeneration [26–31]. Numerous studies have highlighted the utility of conductive biomaterials and how they can facilitate cell adhesion and proliferation [27,32], organization [33], and differentiation [33–35].

Current methods to simulate endogenous tissue conductivity employ the use of electrically-responsive polymers including polypyrrole (PPy) [36], polyaniline (PANI) [37], and poly(3,4-ethylenedioxythiophene) (PEDOT) [38]. These organic conductive polymers exhibit robust electrical properties while maintaining biocompatibility and chemical tunability [38,39]. PPy in particular has proven to be a promising conductive material for the repair of electrically-responsive tissues [40–43]. However, due to difficulties in material processing, such as poor solubility and mechanical brittleness, the majority of conductive biomaterials present cells with a two-dimensional (2D) environment that does not accurately recapitulate the complexity and heterogeneity of their native (3D) environments [28,38]. Furthermore, conductive polymers alone do not possess the anisotropy and ECM cues (e.g., cell adhesive sites) that are necessary for skeletal muscle repair. This difficulty in processing conductive materials into the 3D aligned structures needed for musculoskeletal tissue engineering has proven to be a limitation in the field. A potential solution is the use of hybrid materials composed of conductive polymers and anisotropic cell-instructive biomaterials to leverage the strengths of both components.

For all these reasons, macroporous scaffold-based biomaterials offer an attractive platform for skeletal muscle tissue engineering. Porous scaffolds can provide a 3D interconnected pore structure with large enough pores to allow cellular penetration and growth as well as effective mass transfer of nutrients and metabolic waste. One method that has proven effective for the production of 3D macroporous scaffolds is freeze-drying [28,44,45]. Specifically, collagen-glycosaminoglycan (CG) scaffolds produced via freeze-drying have previously been applied across a wide range of tissue engineering applications including bone [46], cardiac tissue [47], and peripheral nerves [48], and have also been used clinically in the regeneration of skin [49,50]. An additional benefit of this system is that material properties can be easily controlled by altering processing parameters such as freezing and lyophilization rate as well as precursor suspension composition [16,51,52]. As a result, this system is ideal for the facile introduction of conductive polymers into the bulk suspension prior to lyophilization to produce 3D conductive biomaterial composites. Additionally, we have previously developed a directional freeze-drying approach to make aligned 3D CG scaffolds for tendon tissue engineering that could be leveraged for skeletal muscle applications [53].

Although previous studies have employed the use of conductive polymers and architectural cues to direct cellular growth and differentiation, a biomaterials approach to simultaneously mimic the 3D, hierarchical organization, and inherent conductivity of skeletal muscle has proven to be challenging. The development of a biomaterial platform that pairs architectural anisotropy and electrical conductivity in three dimensions could help fill the clinical void for the treatment of VML injuries and other anisotropic conductive tissues. Here, we designed and characterized a 3D electrically conductive and aligned collagen-based scaffold as a potential platform for skeletal muscle tissue engineering and investigated myoblast growth, organization, and maturation.

## 2. Materials and methods

### 2.1. Polypyrrole (PPy) synthesis

PPy microparticles were synthesized via an oxidation reaction with iron (III) chloride (FeCl_3_). 4 g of pyrrole monomer was reacted with 200 mL of 36 mmol FeCl_3_ using vigorous mixing for 24 h under ambient conditions [32]. The resulting black precipitate was isolated via vacuum filtration and repeatedly washed with deionized (DI) water until the washings were clear. The powder was then dried overnight in a vacuum oven before being sieved through a 325-mesh (45 µm) screen. Fourier-transform infrared (FTIR) spectroscopy was used to confirm the chemical structure of the PPy particles. Particle size was quantified from 5 distinct fields of view for 230 particles using ImageJ’s measure function.

### 2.2. Scaffold fabrication

Collagen-glycosaminoglycan (CG) scaffolds were fabricated by directional lyophilization from a suspension of microfibrillar type I collagen from bovine Achilles tendon (Sigma-Aldrich) and chondroitin sulfate derived from shark cartilage (Sigma-Aldrich) in 0.05 M acetic acid (Fig. 1) [53]. A 1.5 wt% collagen and 0.133 wt% chondroitin sulfate suspension was prepared using a high shear homogenizer (IKA) within a recirculating chiller maintained at 4°C to prevent collagen denaturation. Suspension was stored at 4°C until use. For PPy-containing (CG-PPy) scaffolds the microparticles were incorporated into the collagen suspension via vortexing prior to lyophilization. All scaffolds were fabricated using a VirTis Genesis freeze-dryer (SP Scientific). Scaffolds were fabricated via directional freeze-drying using a custom designed mold composed of a conductive copper base beneath an insulating Teflon block. The thermal mismatch within the mold facilitated directional heat transfer causing elongation of the ice crystals in the longitudinal plane during the freezing process and ultimately resulting in anisotropy of the collagen struts [53]. A freezing temperature of −10°C was used for all scaffolds. Following lyophilization scaffolds were dehydrothermally (DHT) crosslinked at 10^5^°C for 24 hrs.

**Figure 1:**
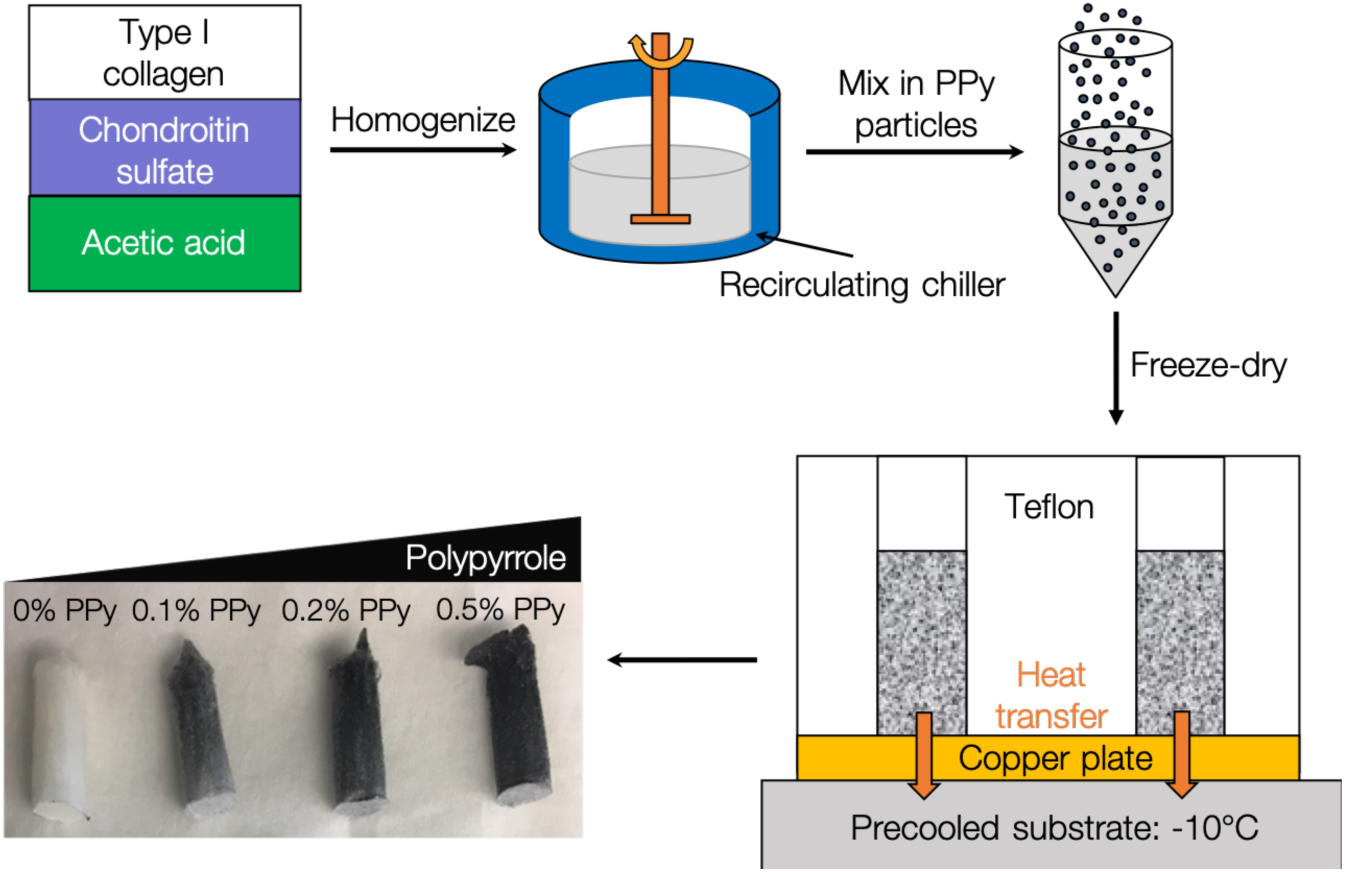
Schematic of collagen-glycosaminoglycan-polypyrrole (CG-PPy) scaffold fabrication. A suspension of type I collagen, chondroitin sulfate, and acetic acid was prepared using a high shear homogenizer in a recirculating chiller to prevent collagen denaturization. Synthesized PPy particles were then added to the slurry and homogenized by vortexing. Aligned scaffolds were produced via directional lyophilization using a custom designed thermally mismatched mold.

### 2.3. SEM analysis

Scanning electron microscopy (SEM) was used to quantify PPy particle size and particle distribution within the CG scaffolds. Additionally, SEM was used to qualitatively assess anisotropy of pore microstructure. SEM analysis was performed with a FEI Quanta 650 scanning electron microscope using a secondary electron detector, backscatter electron detector, and energy-dispersive x-ray spectroscopy (EDS) detector under low vacuum [46]. PPy content was analyzed by tracking chlorine content from the iron chloride dopant. No coating was applied to scaffolds prior to SEM analysis.

### 2.4. Scaffold hydration and crosslinking

Scaffold plugs (7.5 mm diameter, ~ 4 mm thickness) were cut from the middle sections of 15 mm length cylindrical scaffolds. Scaffolds were then sterilized in 70% ethanol for 1-2 h and subsequently washed thrice with PBS. After sterilization scaffolds were crosslinked with using 1-ethyl-3-(-3-dimethylaminopropyl) carbodiimide hydrochloride (EDC) and N-hydroxysulfosuccinimide (NHS) at a molar ratio of 5:2:1 EDC:NHS:COOH where COOH is the carboxylic acid content of the collagen [53,54]. EDC/NHS-mediated crosslinking facilitates the covalent reaction of collagen primary amines with carboxylic acids to improve scaffold mechanical integrity. CG scaffolds were incubated in sterile-filtered EDC/NHS solution for 50 min under moderate shaking before being washed twice with PBS. Scaffolds were stored at 4°C until use.

### 2.5. Pore size analysis

Scaffold pore size (diameter) was analyzed using confocal microscopy across the entire length of the freeze-dried scaffold cylinders to account for any longitudinal differences in pore structure. Scaffold sections were cut to ~ 4 mm thickness using a razor blade. For imaging, scaffolds were stained with a 2 µM solution of AlexaFluor 568 NHS ester in PBS that conjugates to free amines on the collagen scaffold backbone for 20 min under moderate shaking. After staining, scaffolds were washed twice with PBS before imaging. Both transverse and longitudinal scaffold cross-sections were imaged using a Leica SP5 confocal microscope at an excitation wavelength of 568 nm using a 20x magnification objective. Three 10 µm thick z-stacks (1 µm per slice) were taken of each sample and a maximum projection image was created using ImageJ. Image analysis was performed via a linear intercept method using a custom MATLAB program to determine average pore size [55]. Scaffold pore size for both transverse and longitudinal planes was determined from 10 individual images from each scaffold section. 15 scaffolds per experimental group were tested.

### 2.6. Conductivity measurements

Scaffolds used for conductivity measurements were hydrated and crosslinked using the same methodology described earlier. However, all wash steps and chemical crosslinking were performed in DI water to avoid the confounding effects of ions in PBS.

The conductivity of the scaffolds was measured using a platinum electrode parallel plate cell. The length (*L*) of the scaffold was first measured using calipers. Crosslinked scaffolds were blotted to remove excess DI water and loaded into the parallel plate cell. The distance between the electrode plates was then adjusted to ensure uniform contact across the scaffold. EC-Lab® software from Biologic Science Instruments was then used to measure scaffold conductivity using chronopotentiometry. The rate of change of electrical potential within the parallel plate electrode was measured by altering current. The resistance (*R*) of the scaffold was calculated using the slope of the linear regression fit to the chronopotentiometry data using Ohm’s law. Conductivity (*δ*) was finally calculated by Pouillet’s law equation:

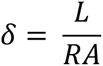

where *L* is length, *D* is diameter, and *A* is the surface area of samples, *A* = *πD*^2^/4 [56]. The conductivity of the scaffolds was modulated by incorporating varying amounts of PPy powder into the collagen suspension. Three scaffolds per experimental group were tested.

### 2.7. Cell culture

Immortalized mouse myoblasts (C2C12s) were acquired from ATCC and used at passages 4 and 5. C2C12s were cultured in standard tissue culture flasks in proliferation media composed of high glucose Dulbecco’s Modified Eagle’s medium (DMEM) supplemented with 10 v/v% fetal bovine serum (FBS, Gibco), and 1 v/v% penicillin streptomycin (Invitrogen). Media was changed every 3 days. For differentiation experiments, C2C12s were cultured in media composed of high glucose DMEM supplemented with 2 v/v% horse serum (Gibco) and 1 v/v% penicillin streptomycin. All cultures were performed at 37°C and 5% CO_2_.

### 2.8. Scaffold culture conditions

C2C12s were seeded and cultured within CG scaffolds containing 0, 0.1, 0.2, or 0.5 wt% PPy. Hydrated and crosslinked scaffolds were incubated in proliferation media for 30 min prior to cell seeding to allow for media adsorption. Cultured C2C12s were trypsinized and resuspended at a concentration of 1.25 × 10^5^ cells per 20 µL media. 10 µL of cell suspension (6.25 × 10^4^ cells) was added to each scaffold and incubated at 37°C for 15 min. After incubation, scaffolds were turned over and seeded with an additional 10 µL of cell suspension for a total of 1.25 × 10^5^ cells per scaffold. Scaffolds were then incubated at 37°C and 5% CO_2_ with proliferation or differentiation media that was changed every 3 days. For differentiation experiments, C2C12s were cultured in proliferation media for 4 days before being transferred to differentiation media.

### 2.9. Metabolic activity

Cell mitochondrial metabolic activity within CG scaffolds was determined using a non-destructive AlamarBlue assay. Viable cells within the construct metabolize the active ingredient in AlamarBlue (resazurin) into a fluorescent byproduct (resorufin) that can be quantified using a fluorescent spectrophotometer [53]. Cell-seeded scaffolds were incubated in AlamarBlue solution for 1.25 h at 37°C with gentle shaking. Resorufin fluorescence was assessed via a Synergy 4 BioTek fluorescence spectrophotometer. The relative cell metabolic activity was determined by interpolating scaffold fluorescence readings using a standard curve derived from known cell numbers. Four scaffolds were tested per experimental group.

### 2.10. Immunocytochemistry

All scaffolds were stained with a 2 µM solution of AlexaFluor 568 or AlexaFluor 633 NHS ester in PBS for 20 min under moderate shaking prior to cell culture in order to visualize the collagen scaffold backbone. After the cell culture period, all scaffolds were fixed in 10% formalin for 30 min prior to further staining.

For cell cytoskeletal visualization, scaffolds were cut in half with a razor blade and the cell membranes were permeabilized with 0.1% Triton X-100 in PBS for 30 min. Scaffolds were washed thrice with PBS and incubated with fluorescein phalloidin (1:200 dilution) in PBS for 30 min to stain F-actin.

For primary antibody staining, following fixation scaffolds were washed with 10 mM glycine in PBS for 5 min and cut in half with a razor blade. Cells were then permeabilized with 0.5% Triton-X-100 in PBS for 30 min followed by another three washes for 5 min with 10 mM glycine. Scaffolds were then incubated for 30 min in protein block solution (Abcam) to block non-specific binding. Scaffolds were incubated overnight with antibodies against myosin heavy chain 4 (MF20, mouse monoclonal, eBioscience, 1:200) or MyoD (mouse monoclonal, Santa Cruz, 1:200) in antibody diluent (BD Biosciences). Next, samples were washed thrice with PBS for 5 min and incubated for 2 h with AlexaFluor 488-conjugated goat anti-mouse (1:200) or AlexaFluor 555-conjugated goat anti-mouse (1:200) secondary antibodies. Subsequently, scaffolds were washed thrice with PBS.

Finally, all scaffolds were stained with a nuclear stain, DAPI (1:1000 dilution) in PBS for 5 min before washing once with PBS. Samples were stored in PBS at 4°C and protected from light until imaging.

### 2.11. Image acquisition

Fixed and stained scaffolds were imaged using a Leica SP5 confocal microscope equipped with Argon-ion, 405 diode (excitation at 488 nm), and white light laser lines (excitation at 568 or 633 nm) to visualize DAPI, F-actin or secondary antibodies, and collagen channels respectively. A 20×0.7 NA objective was used for image acquisition across all samples. Furthermore, the z-stack capabilities of the microscope were used to image through scaffold sections and produce representative images of cell growth and maturation. A PMT detector was used for visualization of the DAPI fluorophore while hybrid detectors were used to image the collagen backbone, F-actin cytoskeleton, and antibody-based stains.

### 2.12. Assessment of scaffold strut and cell alignment

Scaffold pore, cell cytoskeletal, and myotube organization were assessed using the OrientationJ plugin for ImageJ. Collagen strut alignment was determined from 30x SEM images using the ‘Distribution’ function within OrientationJ. C2C12 cytoskeletal alignment was measured after 7 days of culture from 25 µm thick maximum projection images of z-stacks collected using a Leica SP5 confocal microscope. Myotube organization was also assessed following 5 days of culture in differentiation media from 40 µm thick maximum projection images of z-stacks collected using the Leica SP5. For all analysis, images were taken of both transverse and longitudinal scaffold orientations. Scaffold strut and cell alignment is reported in terms of orientation angle (−90°-+90°) for samples taken from the transverse and longitudinal planes [53]. 0° corresponds to the angle of directional solidification in longitudinal sections.

### 2.13. Assessment of cell maturation

The number of nuclei in myosin heavy chain (MHC)-positive cells was determined from twelve different fields of view chosen at random across three separate scaffolds per experimental group. First, the number of nuclei in MHC-positive cells was counted using the binary feature extractor tool in the BioVoxxel toolbox. The filtered nuclei were then used to quantify the number nuclei per myotube using the speckle inspector tool in the BioVoxxel toolbox. The results were expressed as the percentages of MHC-positive cells and the fraction of myotubes containing 1, 2–4, or 5 or greater nuclei.

To measure the myotube diameter, the same twelve fields of view from the MHC analysis described above were used. The DiameterJ ImageJ plugin was used to quantify myotube diameter from maximum projection images of ~ 40 µm thick z-stacks.

In order to evaluate the number of MyoD positive cells, twelve different fields of view were chosen at random across three separate scaffolds per experimental group. The total number of cells was quantified by counting DAPI positive cells using the ‘Analyze Particles’ plugin in ImageJ. An image mask was then made using the DAPI channel and overlaid onto the MyoD channel. Subsequently, the number of MyoD positive cells was determined using the Analyze Particles plugin.

### 2.14. Statistical analysis

Statistical testing was performed using one-way or two-way ANOVAs with Tukey HSD tests using SPSS and R. *P* values < 0.05 were considered statistically significant. Cell metabolic activity data were for *n* = 4 scaffolds per group while scaffold conductivity, pore orientation, and immunocytochemical analyses were for *n* = 3 scaffolds per group. Pore size analyses were for *n* = 15 scaffolds per group. Box plots cover the second and third data quartiles with error bars covering the first and fourth quartiles. Box plots also include marks for mean (*x*) and median (*bar*) data values. Bar graph heights correspond to the mean with standard deviation error bars and individual data points included as scatter plots overlaying the bars.

## 3. Results

### 3.1. Synthesis and characterization of polypyrrole particles

Toward the creation of conductive collagen scaffolds, we first synthesized PPy particles via an oxidation reaction with iron (III) chloride (FeCl_3_) to yield a fine black powder that could be incorporated into CG suspension prior to scaffold fabrication via freeze-drying. Microscopic and spectroscopic characterization was carried out to determine PPy particle physical and chemical properties. SEM images of PPy particles revealed consistent particle size across samples with an average size of 527.1 ± 96.7 nm (**Fig. S1**). PPy chemical structure was also analyzed using FTIR. The peak intensity at 1580 cm^−1^ is associated with the C=C stretching that occurs in the π-conjugated polymer backbone. Additionally, peaks at 1350 and 1220 cm^−1^ are indicative of C-H wagging vibrations and conjugated C-N in-plane stretching, indicating successful synthesis of PPy [57–59].

### 3.2. Scaffold microstructural analysis

After verifying successful PPy synthesis, particles were vortexed into a slurry of type I collagen and chondroitin sulfate in acetic acid and then lyophilized. CG scaffolds were fabricated via directional solidification with a range of PPy content including 0, 0.1, 0.2, and 0.5 wt%. Scaffold pore microstructure was analyzed using SEM. Scaffold anisotropy was then assessed using OrientationJ, an ImageJ plugin. Orientation analysis of scaffold struts showed anisotropic alignment of collagen in the direction of heat transfer during lyophilization (Fig. 2). Collagen struts showed increased frequency of alignment at 0° (denoting vertical alignment) in the longitudinal plane compared to more randomized fiber organization in the transverse plane. Additionally, orientation analysis showed that the incorporation of PPy particles did not affect the open pore microstructure and resulted in similar pore alignment for all experimental groups.

**Figure 2:**
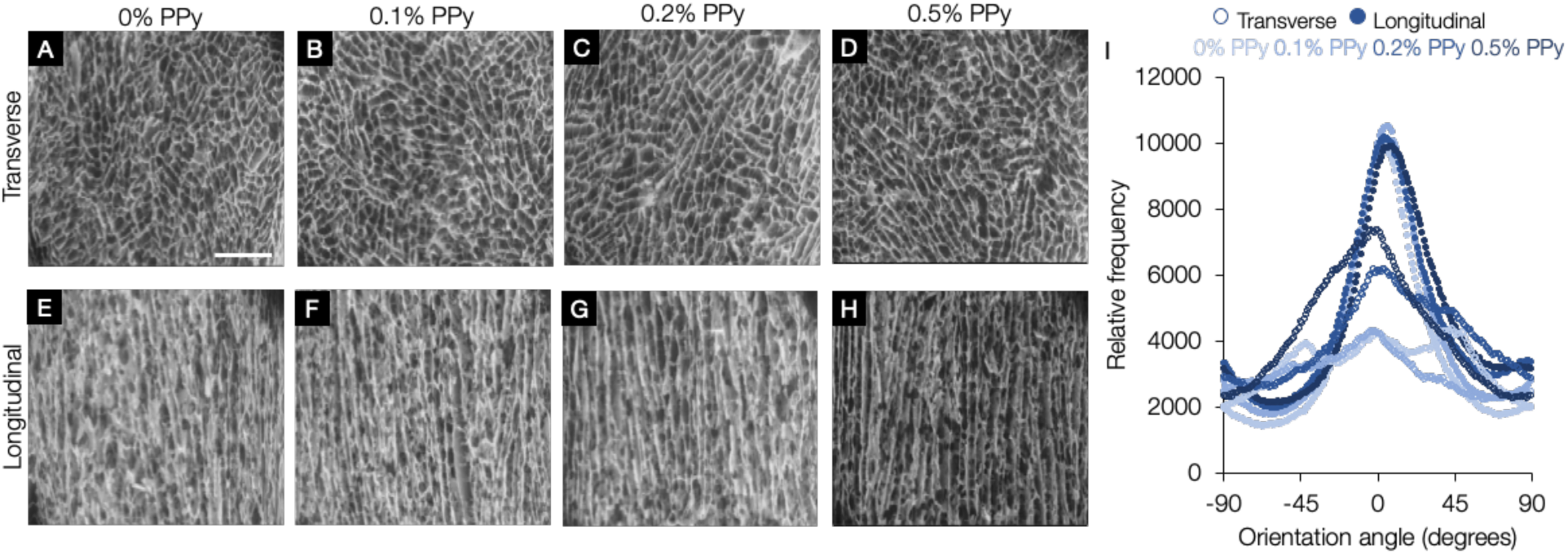
Scaffolds show longitudinally aligned pore orientation independent of PPy content. SEM images (A-D) show isotropic pore structure along the scaffold transverse plane and highly aligned, elongated pores along the scaffold longitudinal plane (E-H) mimicking native skeletal muscle tissue organization. I) Histogram of scaffold strut orientation angles within the transverse (*open circles*) and longitudinal (*closed circles*) planes of the scaffolds. *Scale bar:* 1 mm. *n* = 3 scaffolds per experimental group.

### 3.3. Assessment of polypyrrole distribution within scaffolds

While the addition of PPy did not affect scaffold architecture we also aimed to assess how well distributed the PPy particles were within CG scaffolds using SEM analysis coupled with EDS mapping. Since PPy particles were synthesized using an oxidation reaction with FeCl_3_, the presence of chloride (Cl) dopant within the scaffolds was indicative of PPy localization. EDS mapping showed that PPy was homogenously distributed throughout the CG scaffolds (Fig. 3) as designated by the pink pixels corresponding to Cl content. Additionally, the intensity plot shows that the magnitude of the Cl peak increases as the loading of PPy increases and that no Cl was detected in control CG scaffolds.

**Figure 3:**
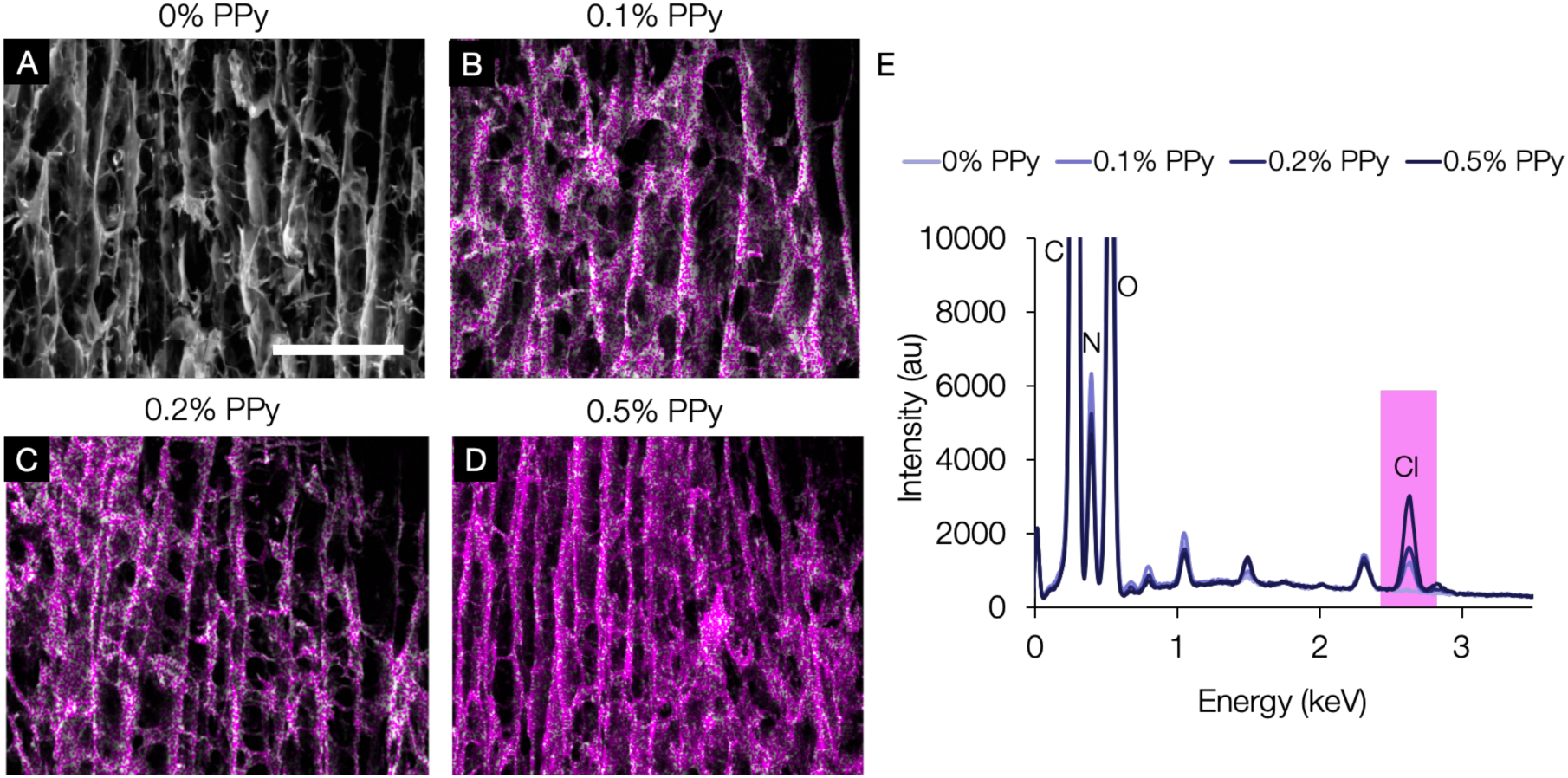
Polypyrrole (PPy) content is uniformly distributed throughout scaffolds. A-D) Energy-dispersive X-ray spectroscopy (EDS) maps of Cl content overlaid on SEM images of longitudinal scaffold planes show increasing Cl content with increasing PPy content as indicated by the pink pixels. Additionally, all maps show uniform Cl distribution and thus PPy particle homogeneity within CG-PPy scaffolds. E) EDS spectra of the CG-PPy scaffolds show elemental distributions across experimental groups, including increasing Cl peak intensity with increasing PPy content. *Scale bar:* 500 µm.

### 3.4. Scaffold pore size analysis

After confirming that PPy particles could be incorporated homogeneously in CG scaffolds without affecting pore alignment, we queried if the addition of PPy affected pore size and geometry [55]. Pores in the scaffold transverse plane generally displayed a rounded morphology with an average pore diameter of 155 ± 27 µm and an average pore cross-sectional area of 17670 ± 532 µm^2^ (Fig. 4). In contrast, pores in the longitudinal plane showed an elongated, ellipsoidal morphology with a larger average pore diameter of 218 ± 49 µm and average pore cross-sectional area of 31696 ± 1610 µm^2^. Additionally, the incorporation of PPy particles did not significantly affect pore microstructure of lyophilized scaffolds in either the transverse or longitudinal planes.

**Figure 4:**
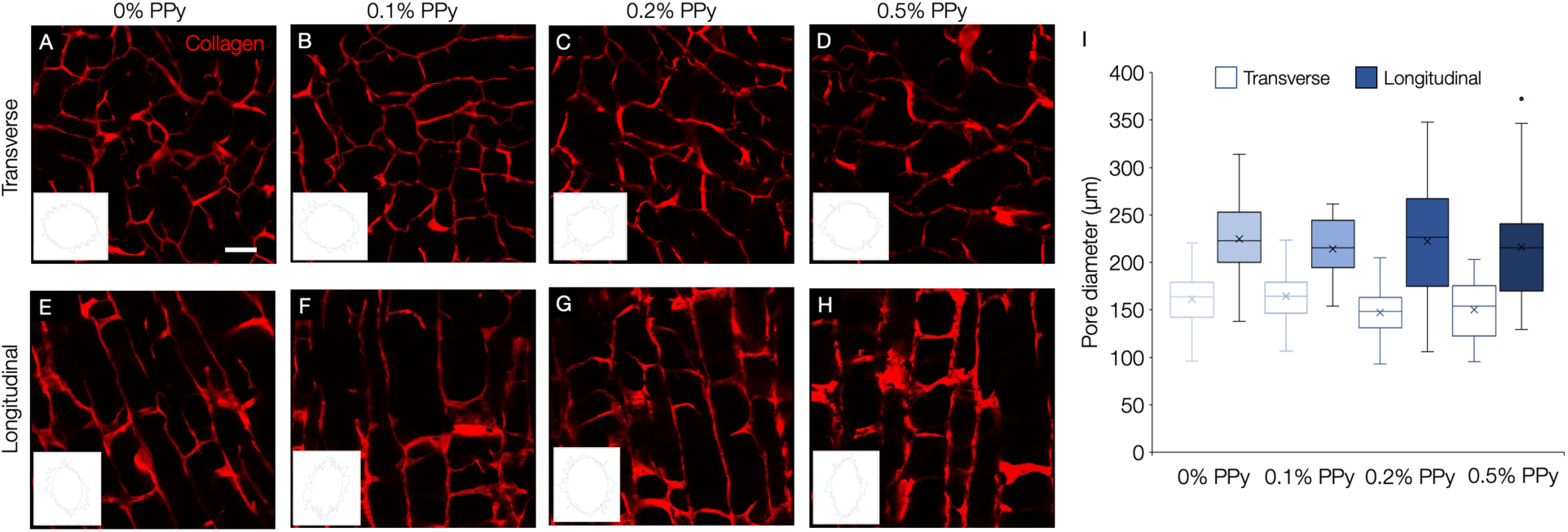
Scaffold pore architecture is not affected by PPy incorporation. Pore microstructure was assessed using MATLAB analysis of confocal images of scaffold collagen backbone. Directional heat transfer during lyophilization resulted in scaffolds containing a 3D pore microstructure with an average pore size of 155 ± 27 µm in the transverse plane (A-D) and 218 ± 49 µm in the longitudinal plane (E-H). *Insets:* best-fit ellipse representations of pore shape generated from MATLAB analysis. I) Box plot of transverse and longitudinal scaffold pore diameters. Data presented for each group includes the mean (*x*) and median (*bar*). Whiskers represent the 1^st^ and 4^th^ quartiles while boxes represent the 2^nd^ and 3^rd^ quartiles. *Scale bar:* 100 µm. *n* = 15 scaffolds per experimental group.

### 3.5. Scaffold conductivity

The conductivity of the CG scaffolds was modulated by incorporating varying amounts of PPy powder into the collagen-chondroitin sulfate suspension prior to lyophilization. The addition of 0.5 wt.% PPy resulted in an approximately five-fold increase in conductivity (1.42 ± 0.18 mS/m) when compared to the collagen scaffold control without PPy (0.27 ± 0.04 mS/m) (Fig. 5) and was significantly greater than all other experimental groups (*P* < 0.01). Moreover, even lower levels of PPy (0.2 wt%) resulted in a significant increase in conductivity compared to the control scaffold with no PPy.

**Figure 5:**
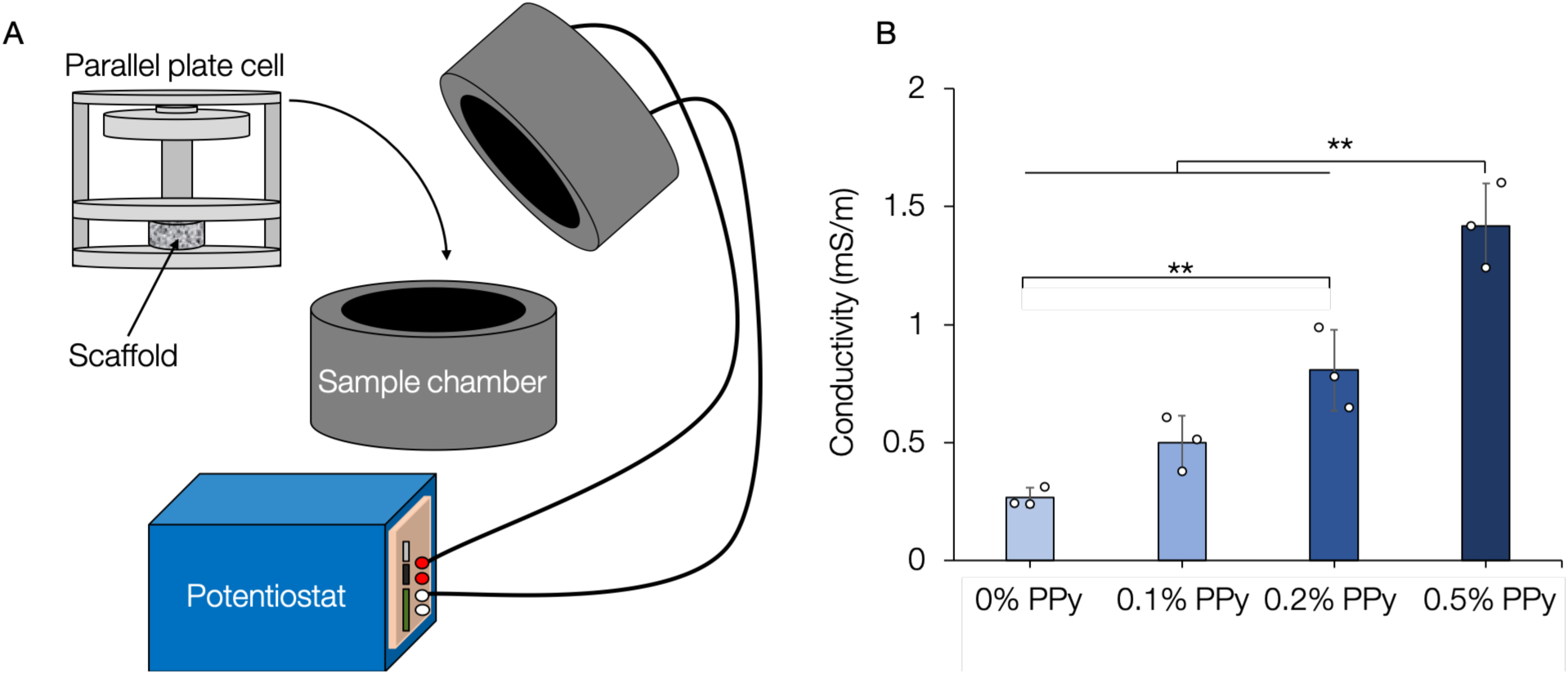
PPy incorporation leads to significantly increased scaffold conductivity. A) Schematic illustrating chronopotentiometry experimental setup. B) The conductivity of the CG-PPy scaffolds was analyzed using chronopotentiometry and indicated that scaffolds with 0.5 wt% polypyrrole content had ~ 5-fold higher conductivity than non PPy-containing control scaffolds. All samples showed increased conductivity with the addition of PPy. **: *P* < 0.01. *n* = 3 scaffolds per experimental group.

### 3.6. Cell metabolic activity in scaffolds

In order to assess how cells would respond to an aligned and conductive 3D environment, immortalized mouse myoblast cells (C2C12s) were cultured within scaffolds for 7 days. Cell mitochondrial metabolic activity, as measured by alamarBlue, indicated that CG scaffolds supported sustained and increasing metabolic activity (Fig. 6) across all experimental groups. While initial cell metabolic activity appeared to be reduced in 0.5 wt% PPy scaffolds after 1 day of culture, metabolic activity was not statistically different between experimental groups after 7 days. These data indicate that the addition of PPy did not detrimentally affect cell mitochondrial metabolic activity. However, preliminary experiments found that increasing the load of PPy in CG scaffolds up to 1 and 1.5 wt% resulted in reduced metabolic activity, indicating that PPy can be cytotoxic at high concentrations (data not shown). As a result, due to its superior conductivity and biocompatibility, the 0.5 wt% PPy-containing CG scaffold was compared to the CG-only scaffold control for all subsequent experiments.

**Figure 6:**
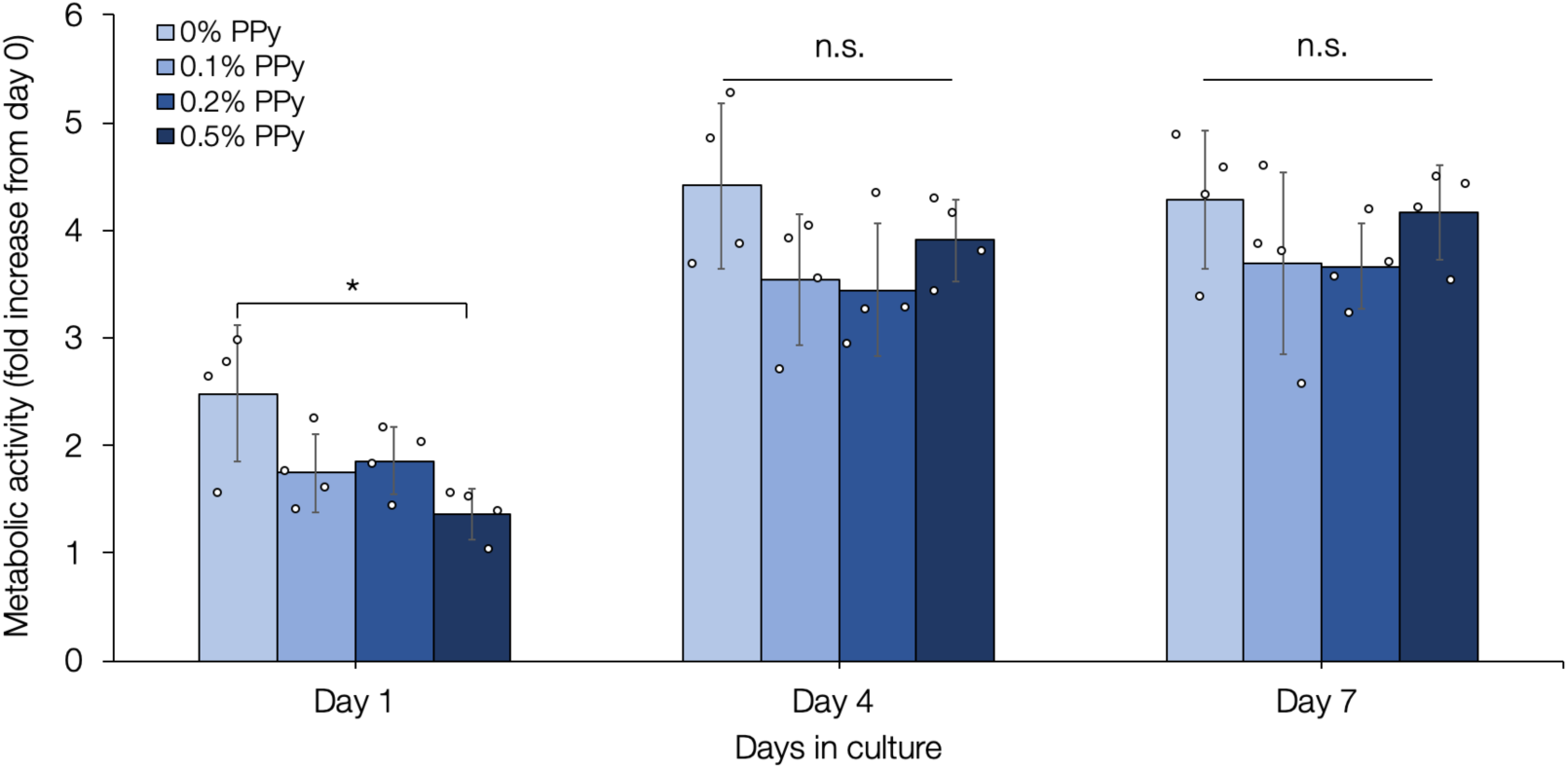
CG-PPy scaffolds support sustained and increasing myoblast metabolic activity. Quantification of metabolic activity over 7 days of culture showed that scaffolds with polypyrrole content ranging from 0-0.5 wt% all supported sustained, increasing metabolic activity. *: *P* < 0.05. *n* = 4 scaffolds per experimental group.

### 3.7. Cell alignment in scaffolds

Following the assessment of myoblast metabolic activity, we aimed to determine if 3D pore alignment could provide contact guidance to organize and align myoblasts cultured within the scaffolds. Confocal images of myoblasts were taken after one week in culture from both the longitudinal and transverse scaffold planes, and were analyzed to determine the relationship between scaffold microstructural cues and cellular alignment. Confocal images indicated that cells conformed to the scaffold contact guidance cues and organized along the collagen backbone in both transverse and longitudinal planes (Fig. 7). In the transverse plane, where scaffolds contained isotropic rounded pores, cell cytoskeletal orientation angles were randomly distributed. However, in the longitudinal plane, myoblasts showed anisotropic cytoskeletal alignment similar to the high degree of organization observed for the CG scaffold backbone. Additionally, myoblast cytoskeletal organization was similar in both 0 and 0.5 wt% PPy scaffolds, indicating that cell organization is dictated by architectural cues.

**Figure 7:**
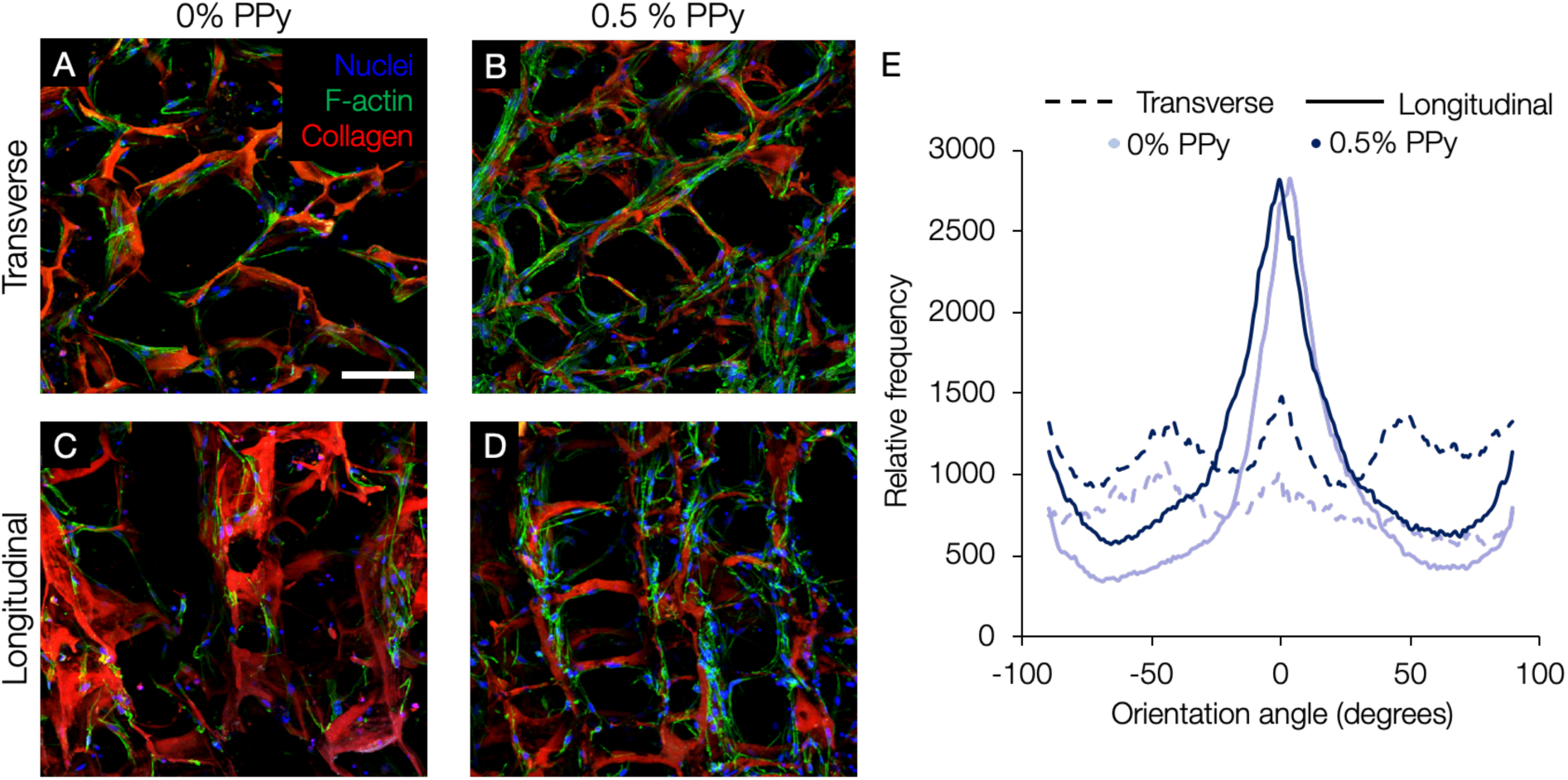
Aligned scaffolds guide 3D myoblast cytoskeletal alignment. Confocal imaging shows that cells spread within CG scaffolds and conformed to contact guidance cues presented by the scaffold microstructure. Cells displayed more random, isotropic organization in the scaffold transverse plane (A-B) and were organized and highly aligned in the scaffold longitudinal plane (C-D). E) Histogram indicating F-actin cytoskeletal anisotropic alignment in the longitudinal (*solid lines*) vs. transverse (*dashed lines*) planes. *Scale bar:* 100 µm. *n* = 3 scaffolds per experimental group.

### 3.8. Determination of myoblast differentiation and myotube maturation in scaffolds

After determining that PPy incorporation did not detrimentally affect cell metabolic activity or inhibit 3D cell alignment within the scaffolds, we aimed to determine if the aligned and conductive scaffold system could facilitate myoblast differentiation. Early C2C12 myogenic differentiation and myotube formation in CG scaffolds were assessed by immunocytochemical analysis of myosin heavy chain (MHC) and MyoD expression after 2 and 5 days in differentiation media (preceded by 4 days of culture in proliferation media after initial cell seeding).

Myoblasts cultured within PPy-containing CG scaffolds generally showed increased MHC staining (**Fig. S2**) corresponding to a higher fraction of MHC-positive cells (33.6% vs 21.7% at day 5, Fig. 8). CG-PPy scaffolds also supported formation of myotubes with more nuclei; by day 5 of differentiation culture 17.1% of myotubes had 5 or more nuclei while only 7.9% of myotubes in non PPy-containing CG scaffolds reached this level (Fig. 8d). Scaffold microarchitecture also influenced myotube organization with myotubes showing preferential alignment in the longitudinal scaffold plane (Fig. 8e). Furthermore, myotube organization was similar between 0 and 0.5 wt% CG-PPy scaffolds, indicating myotube alignment was governed by scaffold contact guidance cues. Although we observed increased MHC signal and presence of multinucleated myotubes in PPy-containing scaffolds, there were no significant differences observed in myotube diameter (**Fig. S3**) or MyoD nuclear localization (**Fig. S4**) between experimental groups.

**Figure 8:**
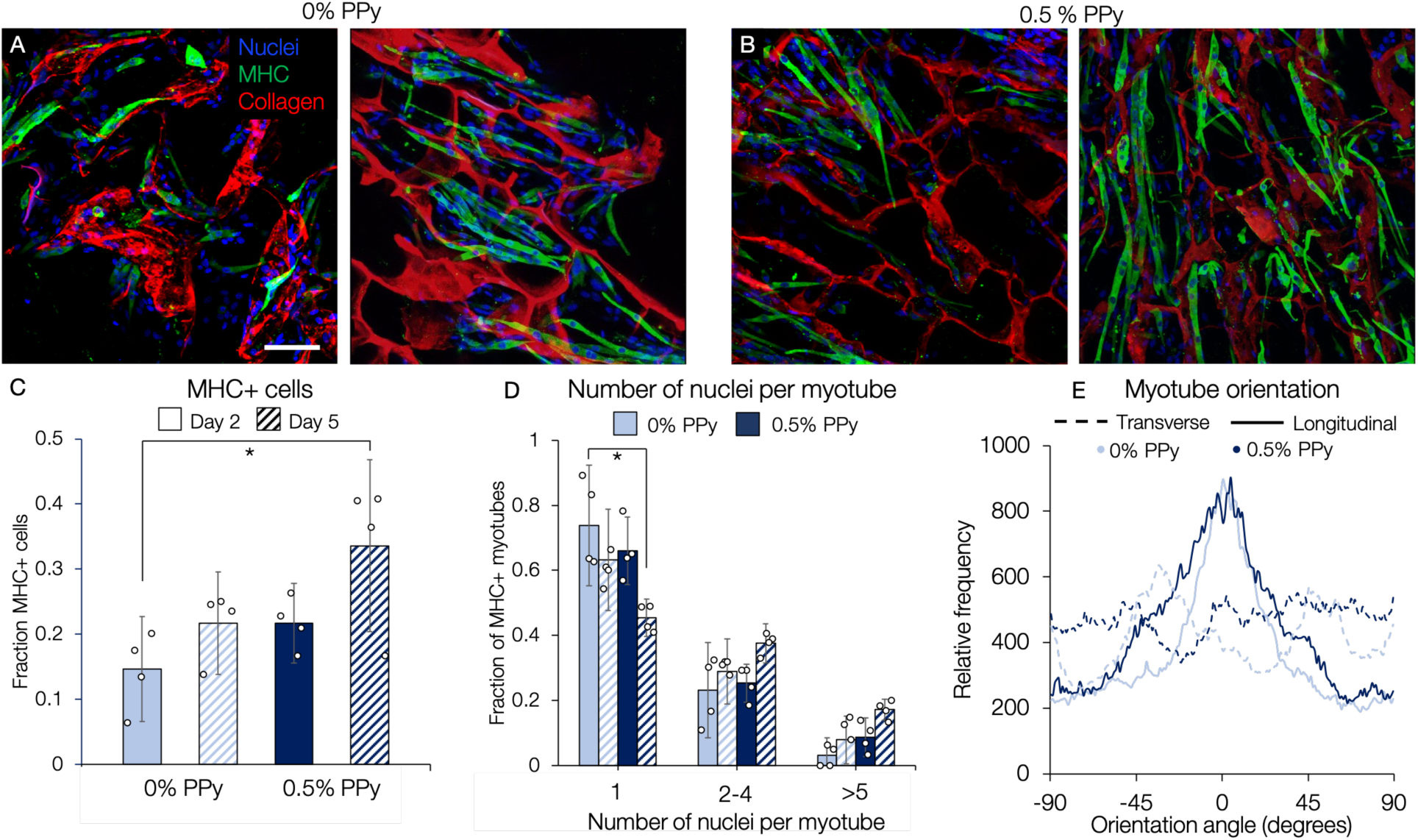
CG-PPy scaffolds support improved early myoblast differentiation. C2C12 myoblasts cultured within CG scaffolds containing A) 0% or B) 0.5% PPy express myosin heavy chain (MHC) after 5 days of culture in myogenic differentiation media. C) PPy-containing scaffolds showed increased fraction of MHC-positive cells after both 2 and 5 days in differentiation media. D) The number of single nuclei MHC-positive myotubes was significantly reduced in PPy-containing scaffolds with higher fractions of multinucleated myotubes observed. E) OrientationJ analysis of myotube alignment indicated preferential alignment in the anisotropic longitudinal plane (*solid lines*) compared to the isotropic transverse plane (*dashed lines*). *Scale bar:* 100 µm. *\**: *P* < 0.05. *n* = 4 scaffolds per experimental group (panels C, D), *n* = 3 scaffolds per experimental group (panel E).

## 4. Discussion

We describe the development of a 3D electrically conductive scaffold with an anisotropic pore structure designed to improve cell growth, alignment, and maturation for skeletal muscle tissue engineering. We chose polypyrrole (PPy) as the conductive component of our scaffold due to its favorable electrical properties, its role in supporting electrically excitable cell behaviors [32,40,41,60], and its minimal immunogenic response following implantation [40,41]. 2D ECM-doped PPy thin films have previously demonstrated the capacity to support C2C12 myoblast adhesion and generation of desmin-positive myotubes [60]. PPy films have also shown the ability to support improved neurite extension of both rat adrenal phaeochromocytoma (PC-12) cells and primary chicken sciatic nerve explants when subjected to electrical stimulation as compared to cells cultured on tissue culture plastic [40]. Importantly, these PPy films elicited minimal adverse tissue response when compared to FDA-approved poly(lactic acid-co-glycolic acid) (PLGA) following either subcutaneous or intramuscular implantation [40].

In this study, PPy particles were chemically synthesized by an oxidizing reaction of pyrrole with iron (III) chloride and added to the collagen-GAG slurry prior to lyophilization. The Cl anion in iron (III) chloride acts as a molecular dopant and renders the PPy conductive. The Cl acts as a charge carrier and when an electrical potential is applied transmits the signal along the π-conjugated polymer backbone [38]. As a result, the Cl shown in the EDS maps is indicative of the presence of PPy and highlights the uniform particle distribution necessary to allow effective charge transfer throughout the scaffold (Fig. 3) [38].

Following scaffold fabrication, conductivity was assessed using chronopotentiometry. All scaffolds were hydrated in DI water and crosslinked using EDC/NHS chemistry, which on its own has been previously been shown to enhance collagen piezoelectric properties [61]. Scaffolds doped with 0.5 wt % PPy were significantly more conductive than all other experimental groups (1.42 ± 0.18 mS/m) (Fig. 5). However, the conductivity of CG-PPy scaffolds is significantly less than the endogenous conductivity of native/mature skeletal muscle (0.708 S/m) [62]. This elevated conductivity of muscle tissue is likely due to interactions between other components of the skeletal muscle niche, specifically extracellular fluid, proteoglycans, and the electrically excitable cells themselves [63]. Additionally, there is currently limited information on the conductivity of the skeletal muscle ECM itself, making comparison with our scaffold material properties difficult. We hypothesized that despite the modest increase in CG-PPy scaffold conductivity, it could still prove an effective mediator of the cell-cell interactions necessary to augment myoblast differentiation, especially given the similarity in magnitude to the mV-scale changes in membrane potential associated with myoblast maturation into myotubes [31,64]. We were particularly encouraged by a recent study that investigated the combined influence of scaffold alignment and electrical conductivity in a 2D electrospun nanofiber system [65]. The authors found that the addition of PANI to anisotropically aligned nanofibers resulted in enhanced C2C12 myoblast fusion and maturation compared to non-conductive and isotropic variants, underscoring the synergy of alignment and conductivity on myoblast differentiation [65]. However, a limitation of the study was that all experiments were confined to 2D constructs that do not mimic the 3D skeletal muscle microenvironment, thus limiting the relevance of the approach to tissue engineering applications.

While previous studies have used freeze-drying to fabricate 3D PEDOT-based conductive constructs for bone [66] and neural [67] tissue engineering applications, we sought to create 3D conductive scaffolds that also incorporated aligned pores to direct cell and ECM organization in a manner mimicking native skeletal muscle. We successfully applied a directional ice templating and lyophilization approach [53,68] to fabricate conductive collagen-GAG-PPy (CG-PPy) scaffolds with a highly aligned macropore structure (Fig. 2). SEM analysis of collagen strut organization showed significant alignment in the longitudinal plane compared to the transverse plane and indicated that the anisotropy was uniform throughout the 15 mm scaffold length. This high level of organization and anisotropy is reminiscent of native/mature skeletal muscle ECM [25]. Moreover, previous work indicated that scaffolds prepared in this manner had a homogenous pore size distribution throughout the scaffold length [53].

Previous work has also used control of freezing kinetics to dictate CG scaffold architecture and demonstrated the importance of pore size on cell attachment and proliferation [69,70]. We chose a freezing temperature of −10°C for scaffold fabrication to promote slower freezing, increased ice crystal coarsening, and formation of larger (150-200 µm) pores. Previous work has shown that scaffolds fabricated at this temperature contain open, interconnected pores that facilitate superior cell adhesion, proliferation, and favorable expression of tissue-specific genes as compared to scaffolds fabricated at lower temperatures with smaller pores [51,53]. This large pore structure (orders of magnitude larger than nanometer-scale mesh sizes in most hydrogels) is also necessary to allow for adequate cell infiltration and migration as well as nutrient transport [28,50,53]. In the case of skeletal muscle tissue engineering, a large pore structure is further advantageous because as myoblasts fuse and mature they increase in size and require more space [23]. We have previously shown the median fiber cross-sectional area of rat tibialis anterior muscle is on the order of 1000 µm^2^ [71], meaning that our scaffold (mean pore cross-sectional area of ~ 20000-30000 µm^2^) should initially support the formation of small muscle fiber bundles. Since CG scaffolds are made of naturally-derived components susceptible to enzymatic degradation over the course of weeks to months (depending on the degree of crosslinking) [72], we predict that gradual cellular remodeling of the scaffold following implantation would permit the formation of larger, more robust muscle fiber bundles.

After characterizing scaffold material properties, mouse myoblasts (C2C12s) were seeded and cultured within CG-PPy scaffolds to assess their capacity to support myoblast adhesion and proliferation. CG scaffolds promoted sustained myoblast mitochondrial metabolic activity over 7 days of culture for PPy concentrations up to 0.5 wt%. However, after 1 day in culture 0.5 wt% PPy scaffolds showed reduced metabolic activity compared to other experimental groups (Fig. 6). This reduction is potentially due to lower initial cell attachment from PPy blocking available binding sites along the collagen scaffold backbone. One potential approach to improve cell attachment would be to modify the PPy with cell adhesive cues; however, additional chemical modifications to the PPy would likely curtail electrical conductivity [32]. Despite reduced initial metabolic activity, myoblast metabolic activity in the 0.5 wt% CG-PPy group increased roughly 3-fold from day 1 to day 4 with no significant differences in metabolic activity found between any experimental groups at days 4 or 7.

After showing that CG-PPy scaffolds could support sustained myoblast metabolic activity, we assessed the ability of CG-PPy scaffolds to facilitate early myoblast differentiation as measured by myosin heavy chain (MHC) expression and myotube formation. The CG-PPy scaffolds supported increased fraction of MHC-positive cells and an increase in number of multinucleated myotubes (Fig. 8). However, many MHC-positive cells remained mononucleated, indicative of an immature phenotype. Although our image analysis approach likely underreported the number of nuclei per myotube by excluding some overlapping myotubes within 3D space, it is likely that studies extending culture times in differentiation media past 5 days would result in more robust myoblast fusion and maturation. An additional limitation of our work with respect to future skeletal muscle tissue engineering applications is the use of an immortalized cell line. Future work will assess primary cell behavior in CG-PPy scaffolds and appraise the efficacy of CG-PPy scaffolds in *in vivo* models of volumetric muscle loss injury.

## 5. Conclusions

We developed a 3D, highly aligned, and electrically conductive collagen scaffold via directional lyophilization of a polypyrrole-doped collagen suspension. This composite biomaterial combines aligned CG scaffolds with conductive PPy particles as a platform for skeletal muscle tissue engineering. Directional lyophilization yielded a highly organized pore microstructure mimicking key features of native skeletal muscle that remained intact after the addition of PPy. EDS mapping and chronopotentiometry confirmed that PPy particles were uniformly distributed throughout lyophilized scaffolds, and furthermore, that increased PPy content resulted in higher conductivity. Directional freeze-drying resulted in a macropore (~ 150-200 µm) structure with elongated pores in the longitudinal plane that was permissive to cell infiltration and early myotube formation. Analysis of myoblast viability indicated that CG-PPy scaffolds supported increasing metabolic activity and that scaffold anisotropy facilitated cytoskeletal organization along the collagen backbone, similar to healthy skeletal muscle. We also showed that PPy-doped scaffolds supported increases in MHC-positive cells and fraction of multinucleated myotubes compared to CG controls. Together, these initial results bode well for the potential application of this material as a scaffold for both *in vitro* studies of cell behavior and *in vivo* guidance of skeletal muscle repair.

## Supporting Information

Characterization of polypyrrole microparticle properties and additional characterization of myoblast differentiation in scaffolds can be found in the supplementary file.

## Data availability

The authors declare that all relevant data supporting this study are available within the paper and the supplementary information, and from the corresponding author upon reasonable request.

## Acknowledgments

The authors would like to acknowledge Dr. Dan Weisgerber and Dr. Brendan Harley for providing the MATLAB code used to assess pore size, Dr. Liheng Cai for use of his confocal microscope, Dr. Geoff Geise, Saringi Agata, and Kevin Chang for assistance with measuring scaffold conductivity, and Dr. Gaurav Giri and Stephanie Guthrie for their help with FTIR. This work was supported by the University of Virginia (UVA), the UVA Biotechnology Training Program (5T32 GM008715), and the NIH (1R21 AR075181). The content is solely the responsibility of the authors and does not necessarily represent the official views of the National Institutes of Health.

