## Supplemental Information for "Aligned and Conductive 3D Collagen Scaffolds for Skeletal Muscle Tissue Engineering"

Ivan M. Basurto[1], Mark T. Mora[1], Gregg M. Gardner[2], George J. Christ[1,3], Steven R. Caliari[1,2*]
[1]Department of Biomedical Engineering, [2]Department of Chemical Engineering, [3]Department of Orthopedic Surgery, University of Virginia

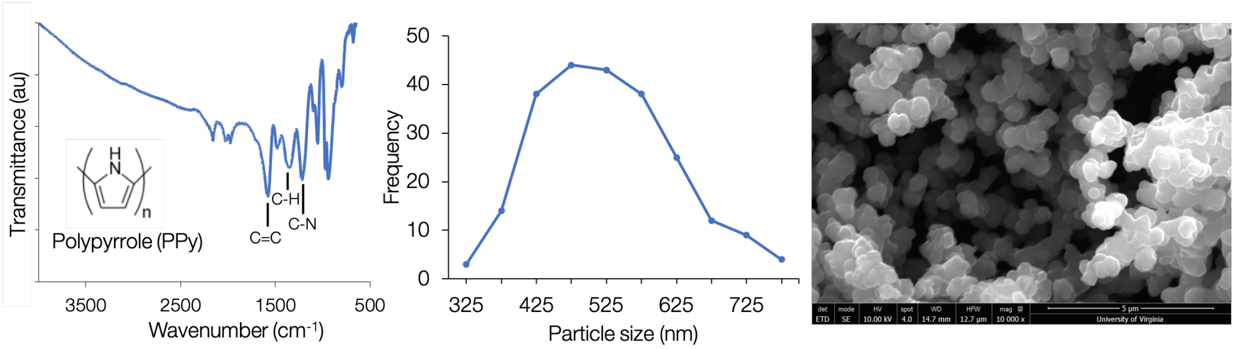


**Supplemental Figure 1.** Polypyrrole microparticles were synthesized via an oxidation reaction and incorporated into a collagen chondroitin sulfate suspension prior to lyophilization. Particle size analysis using scanning electron microscope (SEM) images indicated the production of homogeneous particles with an average diameter of 527.1 ± 96.7 nm. FTIR analysis indicates peaks at 1580 $cm^{-1}$ associated with C=C stretching and peaks at 1350 and 1220 $cm^{-1}$ that are indicative of C-H wagging vibrations and conjugated C-N in-plane stretching. $n$ = 230 PPy particles.

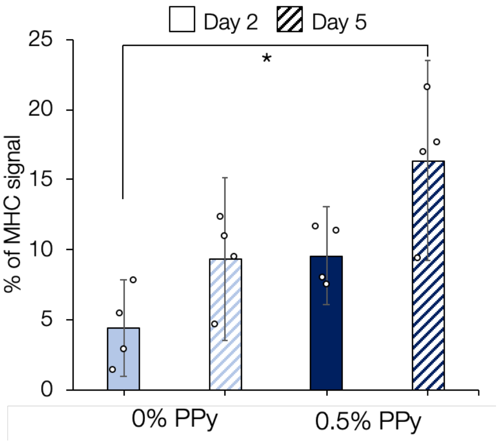


**Supplemental Figure 2.** The percentage of image area occupied by myosin heavy chain (MHC) staining was quantified using ImageJ. After 5 days in differentiation media there was significantly higher signal in the PPy-containing group compared to day 2 signal for the CG-only scaffold control. *: $P < 0.05$. $n = 4$ scaffolds per experimental group.

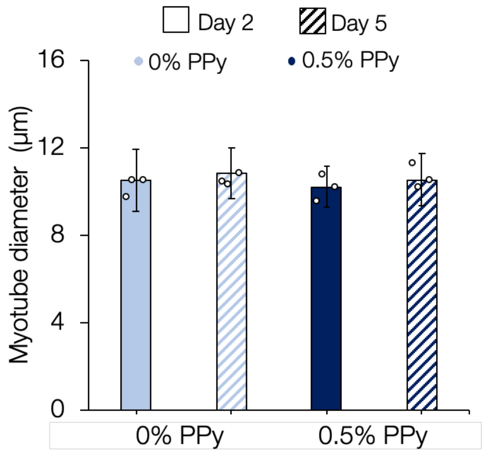


**Supplemental Figure 3.** Myotube diameter was quantified using DiameterJ, an ImageJ plugin. There were no significant changes observed in myotube diameter as a function of culture time or PPy incorporation. $n$ = 3 scaffolds per experimental group.

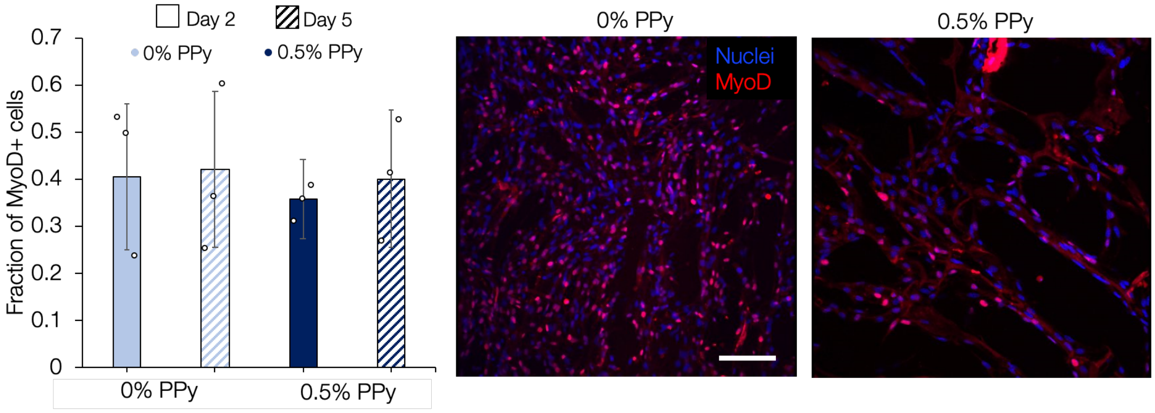


**Supplemental Figure 4.** Assessment of myogenic differentiation indicated that culture time and PPy incorporation did not appreciably affect MyoD expression. *Scale bar:* 100 µm. $n$ = 3 scaffolds per experimental group.
